# Urbanization of a subtropical island (Okinawa, Japan) alters physicochemical characteristics and disrupts microbial community dynamics in nearshore ecosystems

**DOI:** 10.1101/2024.01.12.575464

**Authors:** Margaret Mars Brisbin, Kenneth L. Dudley, Yoshitaka Yonashiro, Satoshi Mitarai, Angela Ares

**Author notes:** College of Marine Science, University of South Florida, St. Petersburg, FL, United States.

## Abstract

Subtropical and tropical islands are undergoing rapid urbanization as human populations and tourism expand worldwide. Urbanization disrupts coastal ecosystems by replacing forests and other natural habitats with roads, parking lots, and buildings. These impervious surfaces increase the amount of runoff and pollution that reaches coastal ecosystems. Urbanization also comes with increased industry, waste treatment needs, fishing and aquaculture pressure, and coastline engineering. Despite the major changes to coasts that accompany urbanization, specific impacts on marine ecosystems can be difficult to measure due to missing baselines. Here, we take advantage of a large gradient in urbanization on the subtropical island of Okinawa, Japan, to evaluate the impact of urbanization on nearshore ecosystems. We measured physicochemical parameters and assessed bacterial community composition every two weeks for one year at two nearshore sites adjacent to watersheds with >70% urban land use and two nearshore sites adjacent to watersheds with >70% rural land use. Our results show that urbanization increases freshwater input and nutrient loading to nearshore ecosystems and profoundly alters the microbial community, overriding the natural seasonal succession observed at rural sites. At urban sites, we detected multiple bacterial species that are fecal indicators and human or marine organism pathogens. The altered physicochemical conditions and microbial communities at urban sites can contribute to the degradation of nearby coral reefs. Results highlight the importance of a “ridge-to-reef” management mindset, as restoring natural coastlines could buffer the impact of urbanization on the marine environment.

## Introduction

Human population density along coastlines is roughly three times the overall global average (Small and Nicholls 2003), and as the world’s human population growth continues, urbanization along coasts is accelerating (Bloom 2011; Seto et al. 2011). The intense concentration of humans in coastal areas disrupts adjacent ecosystems through three main pathways: resource exploitation, pollution, and ocean sprawl (Todd et al. 2019). Resource exploitation includes extraction of living (e.g., fishing, aquaculture) and non-living (e.g., dredging, mining, oil and gas extraction, water use for cooling or desalination) resources (Todd et al. 2019). Pollution near urban centers includes sediments, nutrients, plastic debris, chemicals, and pathogens, as well as increased light and noise (Carlson et al. 2019; Todd et al. 2019). Ocean sprawl includes the development of reclaimed land and artificial islands, artificial coastal defenses, ports, docks, marinas, oil infrastructure, and submarine cables and pipelines (Todd et al. 2019). Each pathway has its own impacts on coastal ecosystems, but they can also act together synergistically, leading to profoundly different ecological conditions near urban centers compared to pre-urbanization conditions. However, absent or shifting baselines make it difficult to fully assess the impact of urbanization on coastal ecosystems (Roberts et al. 2017), as many of the most disturbed ecosystems were impacted by human population centers before ecosystem monitoring was initiated (Röthig et al. 2023).

Like many tropical and subtropical islands with economies fueled primarily by tourism, Okinawa Island (Japan) has undergone urbanization and population growth more recently compared to continental coasts (Uehara et al. 2019; Kobayashi 2022). Okinawa, specifically, experienced rapid urbanization following the 1972 reversion from the U.S.A. to Japanese administration (Li et al. 2018). Okinawa Island is elongated (106 km long by a mean of 11 km wide) and lies on a roughly North-South axis within the Ryukyu archipelago, which demarcates the border between the East China Sea and the Pacific Ocean. Importantly, there is a large population gradient from the south to the north, with sprawling high-density population centers filling the southern third of the island and the northern third of the island being composed of forests, agricultural operations, and small villages. The central third of the island was traditionally agricultural and residential but is quickly giving way to the tourism industry with new hotels developed along beaches and businesses and apartment housing built further inland. Alongside urbanization, Okinawa’s nearshore marine ecosystems have been subjected to resource exploitation (Uehara et al. 2019; Kobayashi 2022), pollution (Li et al. 2018; Ares et al. 2020), and ocean sprawl (Masucci and Reimer 2019)—the three main pathways through which urbanizationi impacts coastal ecosystems (Todd et al. 2019).

Okinawa’s south-to-north ubanization gradient sets up a unique natural laboratory to investigate the ecological outcomes from intense urbanization of a subtropical island and creates the opportunity to fill a critical knowledge gap as islands across tropical and subtropical regions are experiencing increasing tourism demand, growing populations, and accelerating urbanization (Bertolo et al. 2012; Lin et al. 2013). Okinawa also has a strong land-sea connection due to frequent and intense rain events associated with a spring monsoon season and a summer/fall typhoon season (Singh et al. 2022). Okinawa is located in typhoon alley, the most active region for tropical cyclones on Earth (Magee et al. 2021). Consequently, ongoing environmental challenges related to urbanization—particularly stormwater runoff and associated pollutants, such as wastewater and sediment pollution (red soil)—may be especially relevent in Okinawa (Ares et al. 2020).

In this study, we took advantage of Okinawa’s urbanization gradient to investigate how urbanization influences nearshore ecosystems by assessing microbial community composition and physico-chemical environmental parameters every two weeks for one year at four sites with varying levels of urbanization. We focused on microbial community composition because shifting microbial communities indicate or predict large-scale ecosystem disturbances (McLellan et al. 2015). Microbial communities respond rapidly to changing physical and chemical conditions in coastal ecosystems through shifts in both composition and activity. Thus, changes in microbial community dynamics often occur before responses in economically important macro-organisms, such as corals, mollusks, fish, or aquacultured seaweeds. Altered microbial communities are also often correlated with degraded ecosystems (Becker et al. 2023). Moreover, specific microbes are indicators of pollution (e.g., fecal-indicator bacteria) and some microbes are considered pollutants themselves (McLellan et al. 2015). We found that urbanized nearshore ecosystems have significantly different microbial communities and physico-chemical conditions throughout the seasonal cycle when compared to rural ecosystems. We also detected bacteria that are typically associated with wastewater—including several pathogenic groups—at urban sites but not rural sites. Our results highlight the need for comprehensive conservation efforts that include land use management and coastal rehabilitation to ultimately protect important nearshore ecosystems, such as coral reefs and seagrass beds.

## Methods

### Sampling area description

Nearshore seawater samples were collected along the west coast of Okinawa Island in the Ryukyu archipelago, South Japan. The island shows well-defined dominant land uses, with concentrated urbanized watersheds primarily situated in the southern third of the island (Fig. 1). As an exception, Nago City, a major city on the island, lies in the northern part of the island. Four primary sampling sites were selected based on watershed size and the percent of the watershed with land cover classified as urban. Watersheds were delineated and their land areas were calculated using a digital elevation model (DEM) interpolated at 30m based on a 2008 10-meter LiDAR survey provided by the Geospatial Information Authority of Japan (https://fgd.gsi.go.jp/download/menu.php). Land-cover classification was based on a Jan 4, 2015 Landsat 8 Operational Land Imager image obtained from the US Geological Survey (Scene: LC81130422015004LGN00) following methods described in Ross et al. (2018). Land-cover classes were defined as: Agriculture, dominated by sugarcane and other crops at various stages; Forest, dense tree stands with closed canopy; Scrub, short woody vegetation without tree canopy; Grass, short trimmed vegetation found in golf courses and airfields; Rock/Dirt; Sand; Urban, a complex mix of man-made surface materials such as concrete and asphalt with limited vegetation; Water, including fresh and marine bodies of water; and Unclassified, pixels that could not be classified (Fig. 1).

**Fig. 1.**
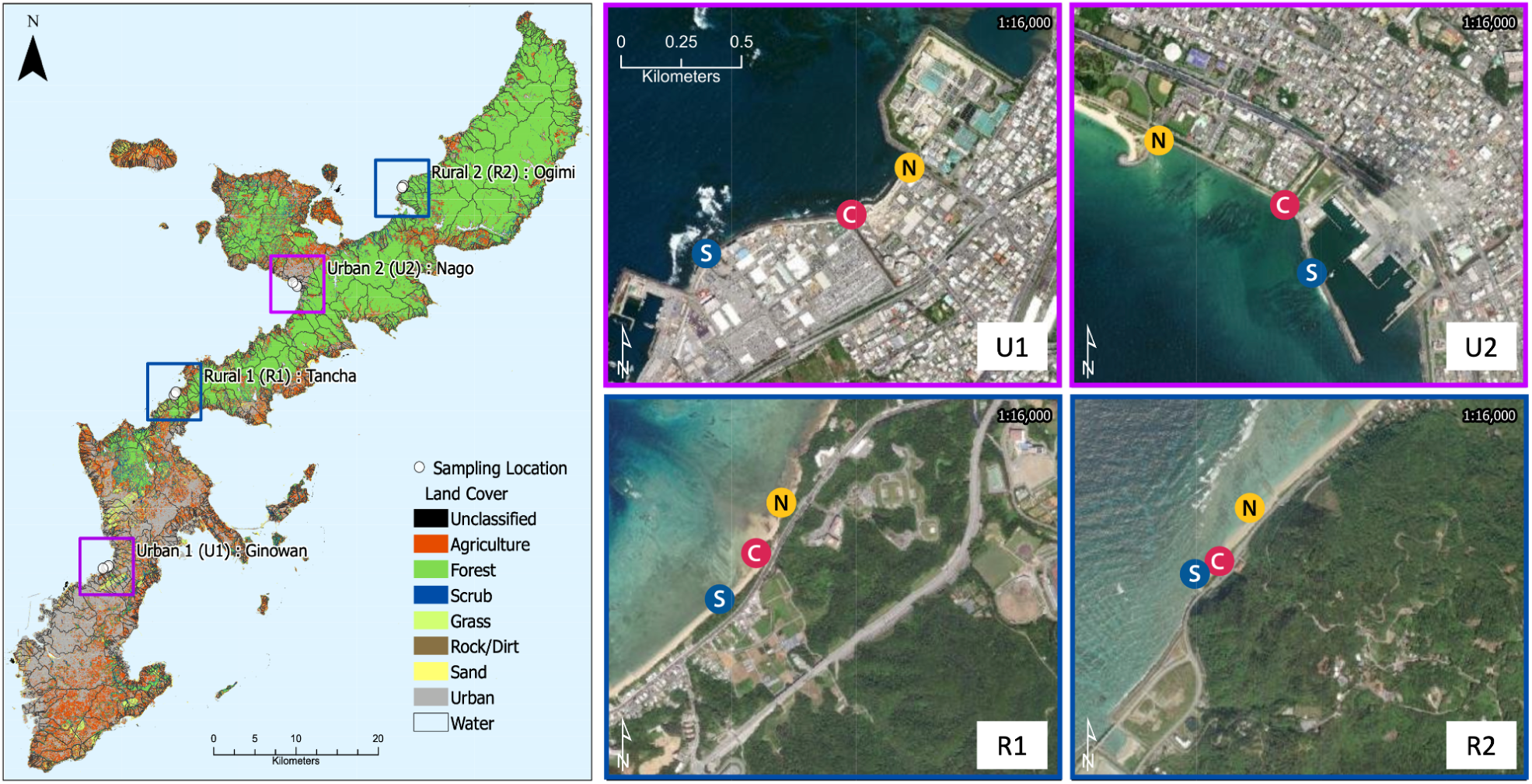
Geographic setting of sampling sites and subsites on the west coast of Okinawa Island, Japan. Left panel: Map of Okinawa Island with land-cover classifications based on satellite data. Thin black lines indicate watershed boundaries based on a LiDAR digital elevation model. Primary sampling sites are highlighted by color coded boxes (purple, urban sites; blue, rural sites) with points demarcating the subsites where samples were collected. Right panels: Detailed maps of the four primary sampling sites with the subsites demarcated as points. Point color indicates the subsites—blue, south (S); red, central (C); yellow, north (N)—where water samples and measurements were taken. Satellite true-color imagery is from Google Earth.

The four primary sampling sites (Urban 1 (U1) - Ginowan, Urban 2 (U2) - Nago, Rural 1 (R1) - Tancha, and Rural 2 (R2) - Ogimi) are located in watersheds of similar size (407,002–993,281 m^2^) and are classified as urban or rural based on the percent of the watershed with land cover classified as urban. Specifically, urban primary sites U1 and U2 are in watersheds with > 70% of land cover classified as urban and rural primary sites R1 and R2 are in watersheds with > 70% land cover classified as agriculture, forest, scrub, grass, rock/dirt, and sand. Primary sampling sites were centered on the freshwater outflow point into the ocean from the study watershed. Three subsites at each primary site were designated for sample collection based on their proximity to the point source of freshwater outflow: central (C) subsites are perpendicularly offshore to the outflow and North (N) and South (S) subsites were 200 m North and South of the outflowss, respectively. We conducted biweekly sample collection for one year, from September 17, 2020–September 2, 2021, leading to 25 sampling events in total.

### Seawater sampling

Nearshore surface seawater was collected for DNA metabarcoding by submerging acid-cleaned 500 mL Nalgene bottles below the sea surface in the uppermost 20–50 cm of the water column. Seawater for dissolved macronutrient analysis was collected in acid-cleaned 50 mL Falcon tubes. A total of 300 of each sample type was collected: four primary sampling sites, three subsites per primary site, and 25 collection events. Samples were transported to the lab on ice and in the dark. After transport, seawater samples for metabarcoding were immediately filtered through 0.2-µm pore-size Polytetrafluoroethylene filters (Millipore) under gentle vacuum pressure, and filters were stored at –80°C for later DNA extraction. Samples for dissolved macronutrient analysis were syringe-filtered and stored at –20°C until chemical analysis. Physicochemical properties—dissolved oxygen (DO), salinity, sea surface temperature (SST), turbidity, and Chlorophyll *a* fluorescence (Chla *a*)—were measured at each subsite with a RINKO conductivity, temperature, and depth (CTD) probe (JFE Advantech, Japan).

### Nutrient analyses

Nutrient concentrations—including Nitrate (NO_3_ ^-^), Nitrite (NO_2_ ^-^), Ammonium (NH_4_ ^+^), Phosphate (PO_4_ ^3-^), and Silica (SiO_2_)—were determined on a QuAAtro39 Continuous Segmented Flow Analyzer (SEAL Analytical) following manufacturer guidelines. Final concentrations were calculated through AACE software (SEAL Analytical). Nutrient Analysis was carried out at the Okinawa Prefecture Fisheries and Ocean Technology Center.

### DNA extraction and metabarcode sequencing

DNA was extracted from frozen filters following the manufacturer’s protocol for the DNeasy Power Water Kit (Qiagen), including the optional heating step. Metabarcode sequencing libraries were prepared for the V3–V4 region of the bacterial 16S ribosomal RNA gene following Illumina’s 16S Metagenomic Sequencing Library Preparation manual without any modifications. Sequencing was performed by the Okinawa Institute of Science and Technology Sequencing Center using 2×300-bp v3 chemistry on the Illumina MiSeq platform. Out of 300 samples, 291 produced sequencing results that passed all quality filters. Overall, 53 million sequencing reads were generated, with 62,656–612,998 sequencing reads per sample (mean = 182,814). Sequencing data are available from the NCBI Sequencing Read Archive (SRA) under the accession PRJNA1044524.

### Bioinformatic and statistical analyses

Sequencing reads were denoised using the Divisive Amplicon Denoising Algorithm (Callahan et al. 2016) with the DADA2 plug-in for QIIME 2 (Bolyen et al. 2019). Taxonomy was assigned to representative amplicon sequence variants (ASVs) using a naive Bayes classifier trained on the SILVA 99% consensus taxonomy (version 132; Quast et al. 2013) with the QIIME 2 feature-classifier plug-in (Bokulich et al. 2018). The results were imported into the R statistical environment (R Core Team 2018) for further analysis with the ‘phyloseq’ (Mcmurdie and Holmes 2013) and ‘vegan’ (Oksanen et al. 2019) R packages. Rarefaction sampling was performed and plotted with the ‘ggrare()’ function and all samples reached richness saturation within their total sample size (Fig. S2). Alpha diversity estimates (richness and Shannon index) were determined with the ‘breakaway’ R package (Willis et al. 2017), and the statistical significance of differences in mean alpha diversity between urban and rural sites each month was tested with pairwise Wilcox tests (Fig. S3, Table S1). To minimize compositional bias inherent in metabarcoding data, we used the Aitchison distance between samples, which includes a centered log-ratio transformation to normalize data (Gloor et al. 2017), for principal coordinate analyses (PCoA). Permutational multivariate analyses of variance (PERMANOVA) on Aitchison distances were performed with the ‘adonis2()’ function (999 permutations) in the R package ‘vegan’ to test whether shifts in community composition between sample types were statistically significant (Oksanen et al. 2019). The influence of environmental parameters in shaping bacterial community composition was investigated through redundancy analysis (RDA) and variance partitioning using functions from the ‘vegan’ R package. Lastly, the ‘SPIEC-EAS’ R package was used for network analysis and to infer co-occurrence patterns between taxa (Kurtz et al. 2015). Intermediate data files and the code necessary to replicate analyses are available in a GitHub repository (https://github.com/maggimars/UrbanOki) where an interactive HTML document (https://maggimars.github.io/UrbanOki/Amplicons.html) can also be found.

## Results

### Land cover effects on physicochemical parameters in nearshore ecosystems

The annual pattern in sea surface temperature (SST) was nearly identical across sites and subsites (Fig. 2). In contrast, salinity was lower and more variable at the urban sites compared to rural sites, especially at the central and north subsites for both urban sampling sites (Fig. 2). Given that watershed size was similar across sampling areas, the lower salinity observed at urban sites is likely caused by more rainwater runoff reaching the coast due to the prevasiveness of impervious surfaces in urban areas. Turbidity was variable at all locations and has multiple causes–including phytoplankton growth, sediment resuspension, and soil pollution (Fig. 2). The percent saturation of dissolved oxygen (DO) was highest at the northern rural site (R2 - Ogimi; all subsites) compared to the other three study sites (Fig. 2), potentially caused by more wave action in that region. Nitrate + nitrite, ammonium, and phosphate concentrations were low at rural sites, whereas these macronutrients were elevated at urban sites—particularly at the central and north subsites of the U1 - Ginowan sampling area (Fig. 3). These results show that urbanization increases macronutrient input in coastal areas. Silica concentrations were variable at all sites (Fig. 3). Silica input may derive from different sources at urban and rural sites—runoff from impervious concrete in urban areas can be high in silica (Maguire and Fulweiler 2016), while runoff including soil in rural areas can also contain large amounts of silica (Ares et al. 2020). Decreases in salinity were strongly correlated with high nitrate + nitrite concentrations at urban sites and were also correlated with increased phosphate and Chl *a* (Fig. 4). Nitrate + nitrite concentrations were also strongly correlated with phosphate concentrations at urban sites and chlorophyll *a* was positively correlated with nitrogen and phosphate at urban sites but not rural sites (Fig. 4). Overall, chlorophyll *a* fluorescence was higher at urban sites compared to rural sites, particularly at central and north subsites (Fig. S1). These results further demonstrate that freshwater input is a source of macronutrients, especially in urban areas, and also suggest that phytoplankton are responding to nutrient input. At rural sites, salinity was significantly negatively correlated with silica concentrations (Fig. 3), indicating that freshwater input is a silica source in these regions.

**Fig. 2.**
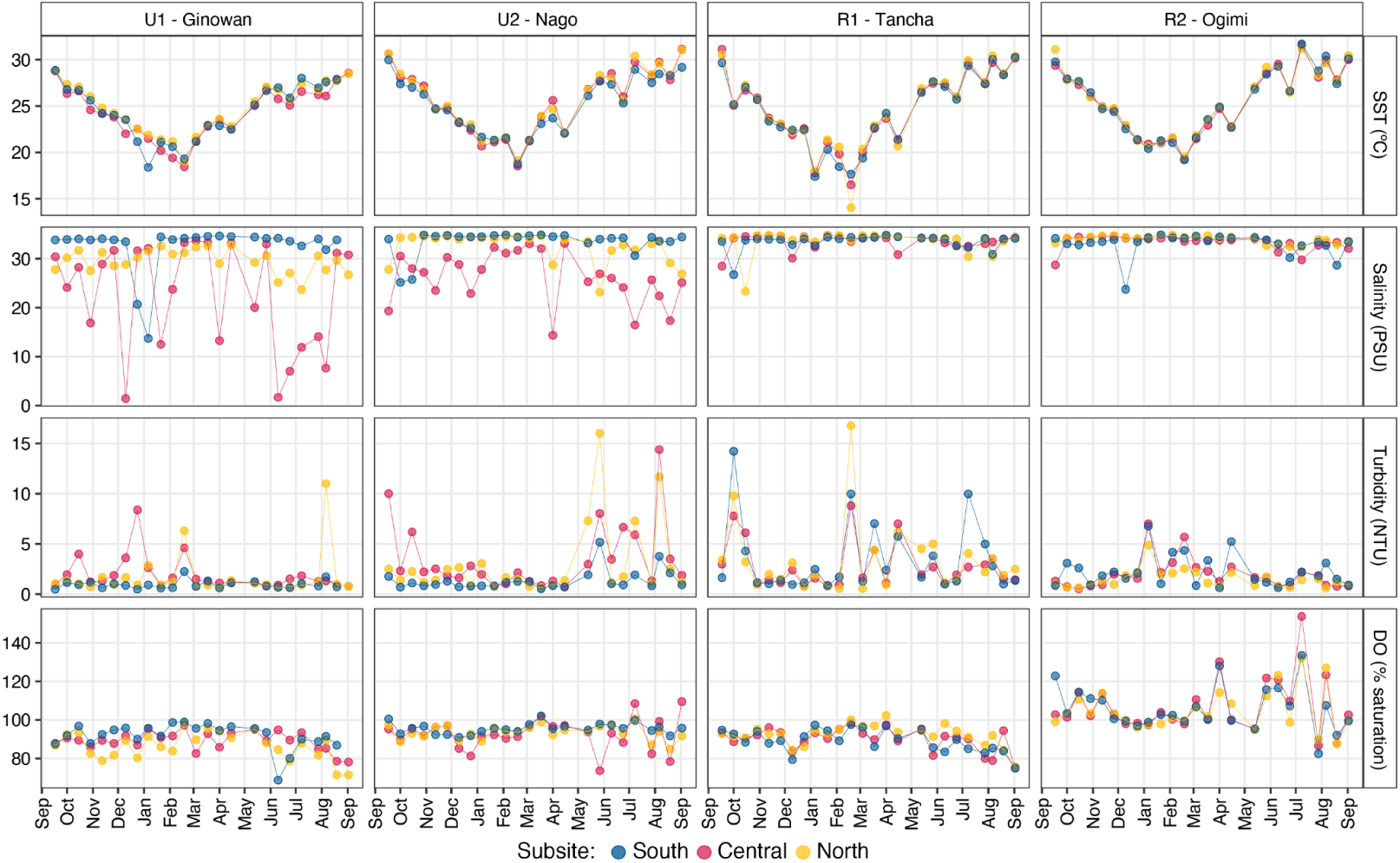
Time series of physical parameters measured at nearshore urban and rural sites along the west coast of Okinawa Island, Japan. Sea surface temperature (SST), salinity, turbidity, and dissolved oxygen (DO) were measured at each subsite (south, central, and north) within each primary sampling site (U1, U2, R1, and R2) using a RINKO CTD probe. Plots are faceted by primary sampling site (vertically) and parameter (horizontally). Point and line color represent subsites—blue, south (S); red, central (C); yellow, north (N)—where water samples and measurements were taken. The annual pattern in SST was nearly identical across sites and subsites. Salinity was lower and more variable at the central and north subsites for both urban sampling sites. Turbidity was variable at all locations. The percent saturation of DO was highest at the northern rural site (R2 - Ogimi)

**Fig. 3.**
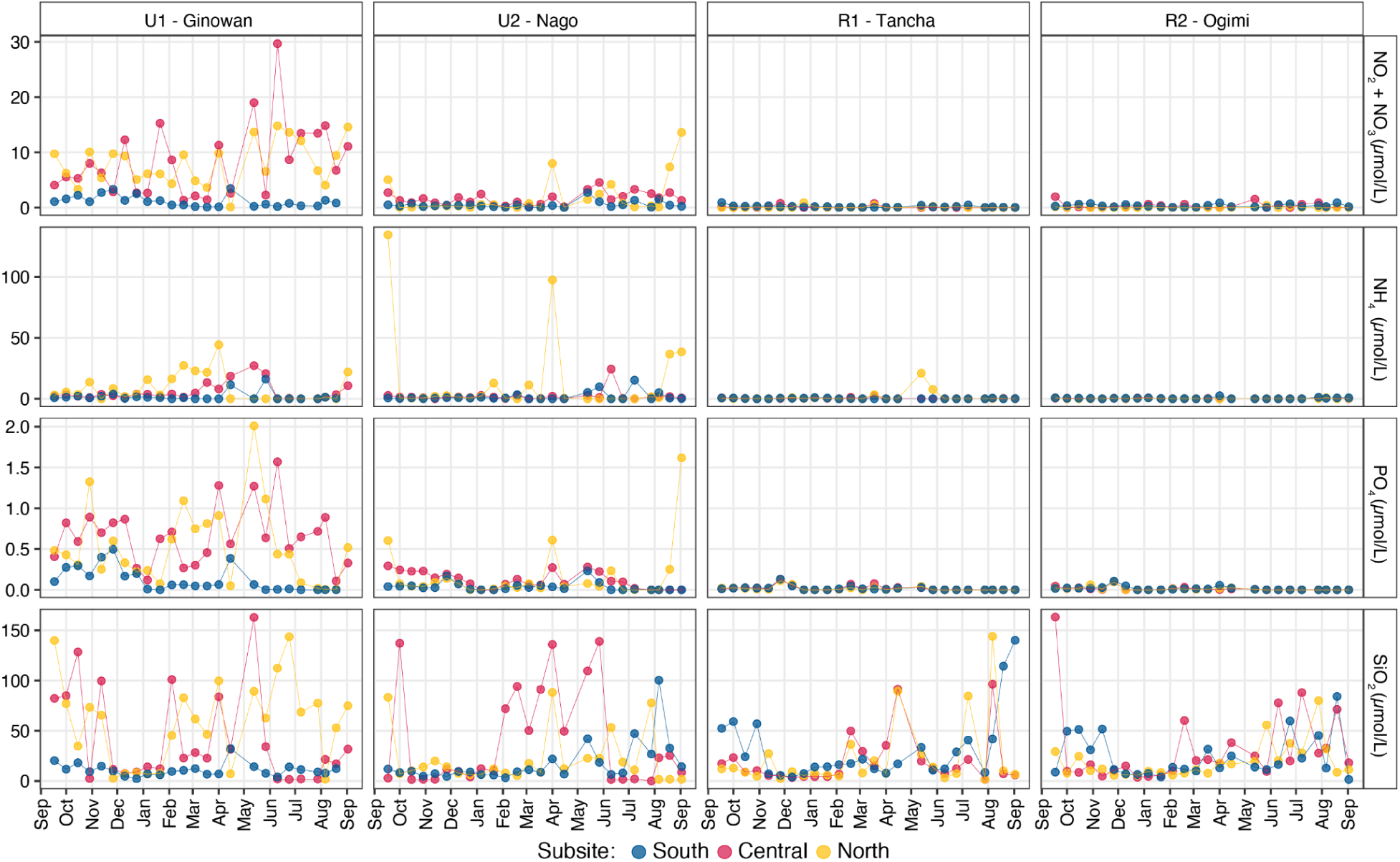
Time series of chemical parameters measured at nearshore urban and rural areas along the west coast of Okinawa Island, Japan. Nitrate + nitrite (NO_2_ + NO_3_), ammonium (NH_4_), phosphate (PO_4_), and silica (SiO_2_) were measured in water samples collected from subsites (south, central, and north) within each primary sampling site (U1, U2, R1, and R2) with a QuAAtro39 Continuous Segmented Flow Analyzer. Plots are faceted by primary sampling site (vertically) and parameter (horizontally). Point and line color represents subsites—blue, south (S); red, central (C); yellow, north (N)—where water samples and measurements were taken. Nitrate + nitrite, ammonium, and phosphate concentrations were rarely above the detection limit at rural sites, while these macronutrients were elevated at urban sites—particularly the central and north subsites of the U1 - Ginowan primary sampling site. In contrast, silica was variable at all sites

**Fig. 4.**
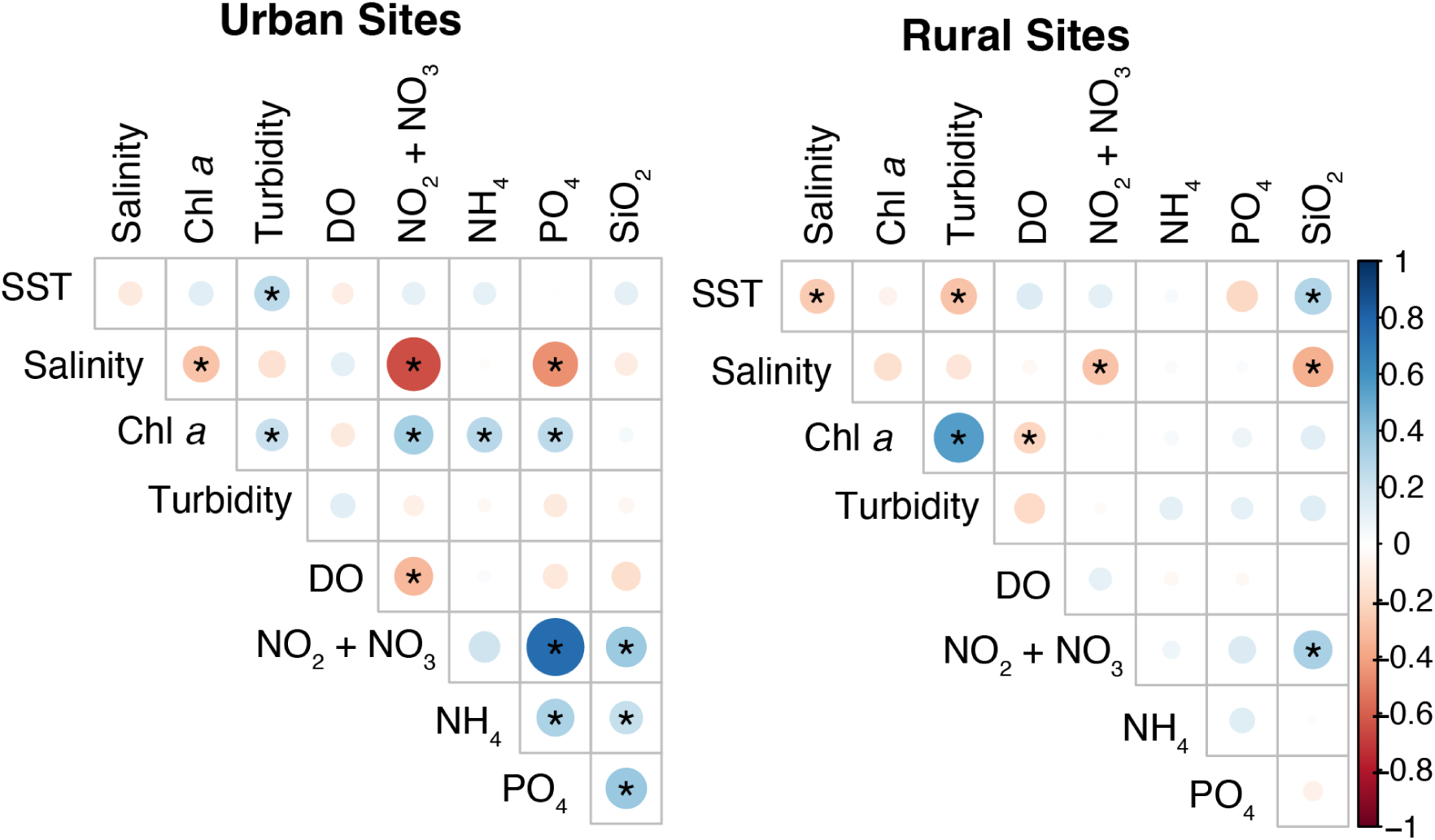
Correlation between environmental parameters at urban and rural sites on the west coast of Okinawa Island, Japan. Values for all parameters were z-scaled and Pearson correlation coefficients were calculated for all parameters at the two urban primary sites (U1 - Ginowan, U2 - Nago) and the two rural primary sites (R1 - Tancha, R2 - Ogimi). Correlation coefficients are visualized for all comparisons by both point size and color: darker blue colors are more positively correlated, darker red colors are more negatively correlated, and point size reflects the absolute value of the correlation coefficient. The statistical significance of each correlation coefficient was evaluated with the ‘cor.mtest()’ function in the ‘corrplot’ R package. Correlation coefficients were considered statistically significant if the *p*-value was ≤ 0.01 and significant correlations are marked with an asterisk (*) on the plot

### Effects of land cover on near-shore bacterial communities

We used metabarcode analysis of the 16S rRNA gene to investigate the effect of land cover (urban or rural) on nearshore bacterial community composition across a biweekly time series spanning a full year. Alpha diversity metrics (ASV richness and Shannon index) were lower in warmer months (May–August) compared to cooler months at both urban and rural sites (Fig. S3). However, the mean richness and Shannon indices were higher at urban sites compared to rural sites across all months (Fig. S3, Table S1). The bacterial community compositions in nearshore waters of urban sites were significantly different from the communities in the nearshore waters of rural sites (PERMANOVA, 999 permutations, *F*=8.5, *p*=0.001; Fig. S4). Principal coordination analysis (PCoA) plots for nearshore bacterial communities in urban and rural regions showed contrasting patterns (Fig. 5). The warmer months (May through the beginning of August) formed tight clusters in the PCoA plots for both rural (Fig. 5a) and urban sites (Fig. 5b). In the PCoA plot for rural sites, samples from seasons cluster so that they are separated roughly by quadrant, with winter samples in the top right, spring and early summer samples in the top left, samples from late summer in the bottom left, and autumn samples in the bottom right. In the PCoA plot for urban sites, samples separate based on which urban site they originated from, with more samples from U1 - Ginowan in the bottom half of the plot and more samples from U2 - Nago in the top half of the plot. These differences in clustering patterns suggest that the seasonal succession cycle is disrupted at urban sites compared to rural sites. However, results from PERMANOVA analyses on the season and primary site variables were significant for both rural and urban sites (Table 1). The subsite variable was only significant for urban sites, indicating that proximity to a fresh water outlet has a larger impact on nearshore bacterial community composition in urban areas than in rural areas.

**Fig. 5.**
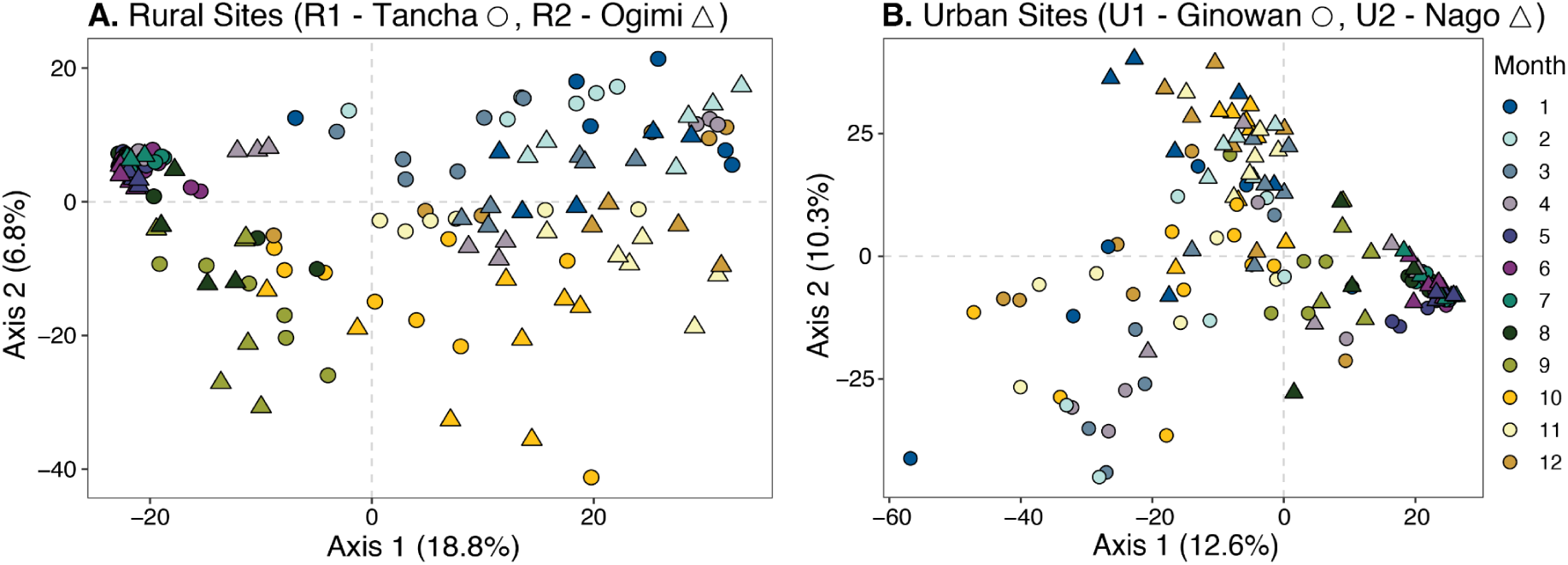
Principal coordinate analysis (PCoA) plots of Aitchison distances between bacterial communities through time at two nearshore sites adjacent to rural areas (A) and two nearshore sites adjacent to urban areas (B). Shape indicates sampling sites, with circles representing the more southern site of both the rural and urban sites, and triangles representing the more northern sites. The color indicates the month of the year, with blues representing winter months, purples representing spring months, greens representing summer, and yellows representing autumn. Samples from late spring and early summer form tight clusters in both plots. Samples from rural sites (A) separate into quadrants based on the season (winter; top right, autumn; bottom right, late summer; bottom left, spring and early summer; top left), whereas samples from the two urban sites (B) separate by site location on the second axis and the seasonal cycle is not visible. A color-blind accessible version of this plot is available at https://maggimars.github.io/UrbanOki/Amplicons.html#with_colorblind_pallette

**Table 1.** Permutational multivariate analyses of variance (PERMANOVA) results for tests run on rural and urban nearshore bacterial communities with month, primary site, and subsite variables.

| Land cover | Variable | *F*-statistic | *R*<sup>2</sup> | *p* |
| --- | --- | --- | --- | --- |
| Rural | Month | 5.7 | 0.32 | 0.001 |
|  | Primary site (R1 v. R2) | 3.2 | 0.02 | 0.002 |
|  | Subsite (S, C, N) | 0.68 | 0.01 | 0.99 |
| Urban | Month | 3.2 | 0.21 | 0.001 |
|  | Primary site (U1 v. U2) | 6.26 | 0.4 | 0.001 |
|  | Subsite (S, C, N) | 3.63 | 0.05 | 0.001 |

We performed a redundancy analysis (RDA) to visualize which environmental variables contributed to the clustering observed in PCoA ordination plots (Fig. S5). An ANOVA was run on the RDA to determine if model results were significant and an ANOVA by term was used to test which variables’ contributions were statistically significant (Table S2). Variance partitioning was calculated for all significant variables. Sea surface temperature (SST) significantly affected community composition clustering among samples from both urban (*F*=3.6, *p*=0.004, variance partition=0.7%) and rural sites (*F*=4.2, *p*=0.012, variance partition=2.2%). Dissolved oxygen (*F*=13.4, *p*=0.001, variance partition=7.4%), phosphate (*F*=6.9, *p*=0.004, variance partition=3.1%), and salinity (*F*=2.83, *p*=0.045, variance partition=0.6%), were also significant determinants of community composition in rural samples. In contrast, nitrate + nitrite (*F*=5.4, *p*=0.001, variance partition=1.5%) was the only significant determinant of community composition in urban samples after SST (Fig. S5, Table S2). The larger influence of phosphate on bacterial communities in rural nearshore waters likely reflects the importance of agricultural fertilizer in runoff in these regions (Hart et al. 2004). Similarly, the larger influence of nitrate and nitrite on bacterial communities in urban inshore waters reflects the importance of nitrogen loading from wastewater and runoff in shaping bacterial communities in these regions (Archana et al. 2016).

By plotting the relative abundance of ASVs grouped by bacterial order, it was evident that common marine bacteria were prevalent across all samples (Fig. 6, Fig. S6). Bacteria in the order Flavobacteriales were the most abundant across all primary sites, subsites, and months (Fig. 6, Fig. S6). Members of this order are among the most abundant picoplankton in the global ocean and can be dominant in open and coastal waters, and in productive and oligotrophic waters (Gómez-Pereira et al. 2010). The SAR11 clade and the orders Rhodobacterales, Synecococcus, and Puniceispirillales (SAR116) were also abundant in almost all our samples. SAR11 bacteria are the most numerically abundant organisms in the global ocean and are especially abundant in warm oligotrophic regions (Morris et al. 2002). Bacteria in the Rhodobacterales order (Dang et al. 2008) are abundant throughout the world’s oceans and are dominant particle-associated bacteria (Dang et al. 2008). Since our samples were not size-fractioned, the Rhodobacterales bacteria we detected could have been particle-associated. Synechococcus is a globally abundant cyanobacteria that is especially abundant in the western Pacific Ocean (Flombaum et al. 2013). Finally, Puniceispirillales (SAR116) are widespread in oligotrophic regions and are a major dimethylsulfide-producing group in the Kuroshio Current near Okinawa (Choi et al. 2015).

**Fig. 6.**
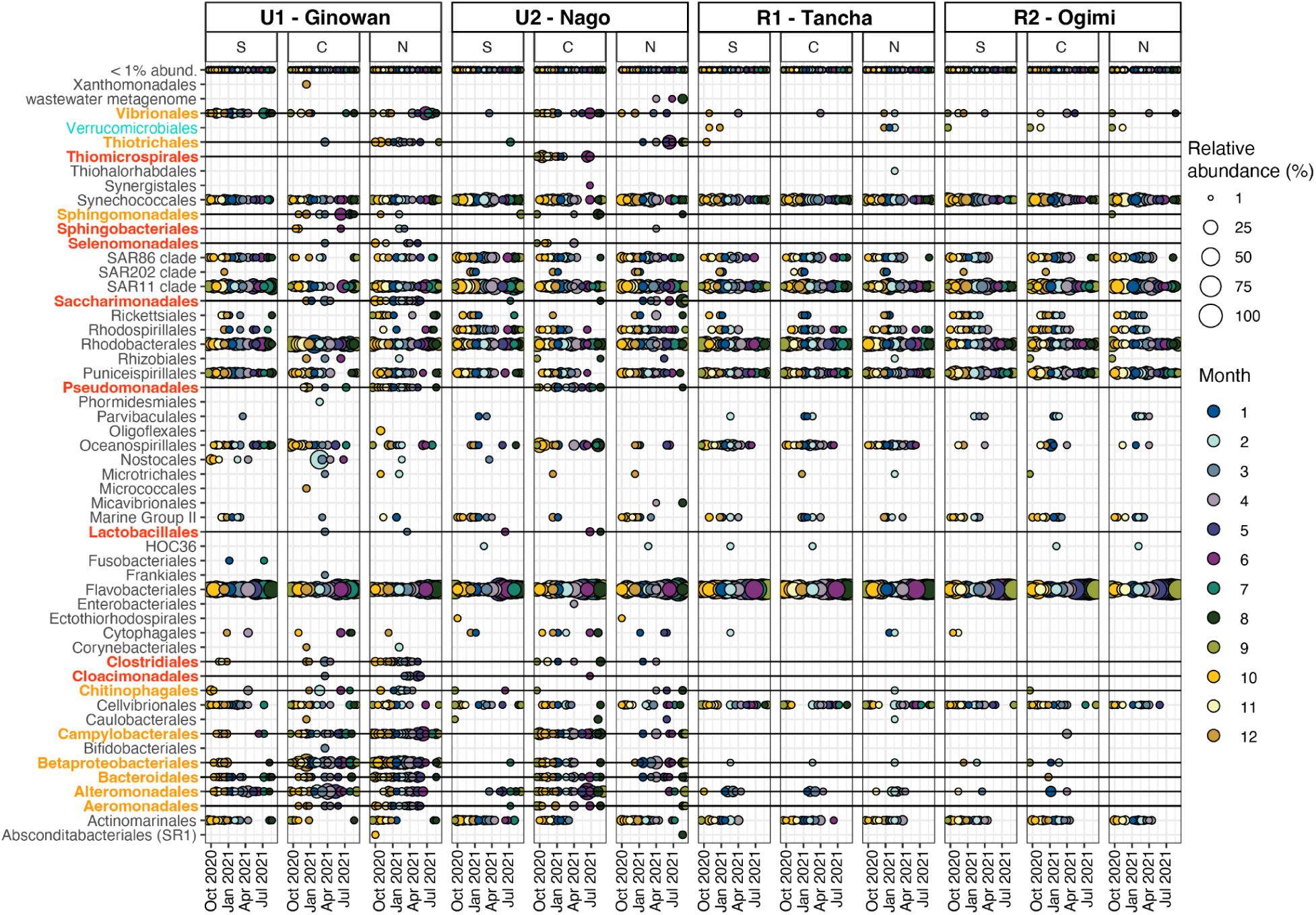
Relative abundance of bacterial orders in each sample. Bubble size represents the relative abundance of each order and was determined by summing the relative abundance of each ASV classified as belonging to the order. Samples are grouped by primary site and columns represent subsites (S; southern subsite, C; central subsite, and N; northern subsite). Bubble fill color indicates the month of the year, with blues representing winter months, purples representing spring months, greens representing summer, and yellows representing autumn. Order names are highlighted as follows: Orange; orders that are prevalent among samples from urban sites but rarely found in rural samples, Red; orders that are prevalent among or present in ≥ 3 samples from urban sites but not found in any rural samples, Green; orders that are prevalent among or present in ≥ 3 samples from rural sites but not found in any urban samples

Aside from the similarities in marine bacterial community compositions across our samples, there was also clear differentiation between samples collected from urban and rural sites. Eight bacterial orders were present in three or more urban samples but absent in all rural samples (Fig. 6; highlighted in red) and nine orders were prevalent among urban samples but rare in rural samples (Fig. 6; highlighted in orange). The cumulative relative abundance of these orders ranges from 1.3–63.7% (mean=19.6%) at subsites in Ginowan (U1) and 1.1–57.9% (mean=15.4%) at subsites in Nago (U2), whereas the cumulative relative abundance of these orders never exceeded 9.8% at rural sites (Fig. 7). Members of these orders were more abundant at the central and north subsites of the two urban sampling sites than at the south subsites (Fig. 7), which are also the subsites that appear more impacted by runoff and freshwater input based on physicochemical parameters (Fig. 2, 3). Interestingly, seven of these orders were shown to covary in urban samples through network analysis (Fig. 8a; Pseudomonadales, Clostridiales, Saccharimonadales, Campylobacterales, Bacteroidales, Betaproteobacteriales, Thiotrichales). The covariance of these orders suggests they may share a common source or that their growth is supported by anthropogenic inputs.

**Fig. 7.**
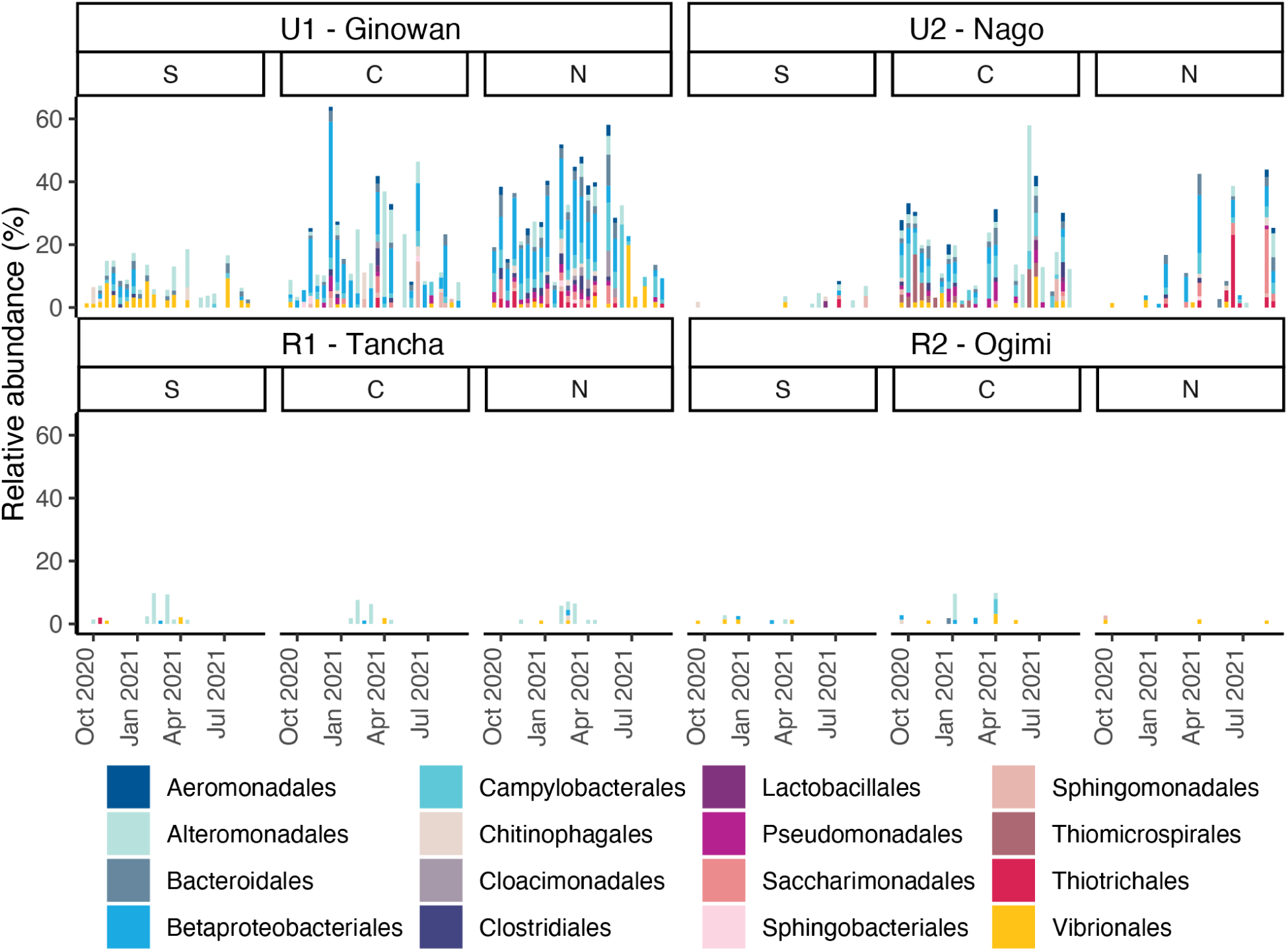
Relative abundance of bacterial orders identified as either prevalent among urban samples and rare in rural samples or prevalent among/present in ≥3 urban samples and absent in all rural samples. Samples are grouped by site and subsite (S; southern subsite, C; central subsite, and N; northern subsite) and stacked bar chart fill colors indicate bacterial order. All orders highlighted in red or orange in Figure 6 are included. Orders that are rare at rural sites make up a large proportion of communities in urban samples, particularly at the central and northern subsites

**Fig. 8.**
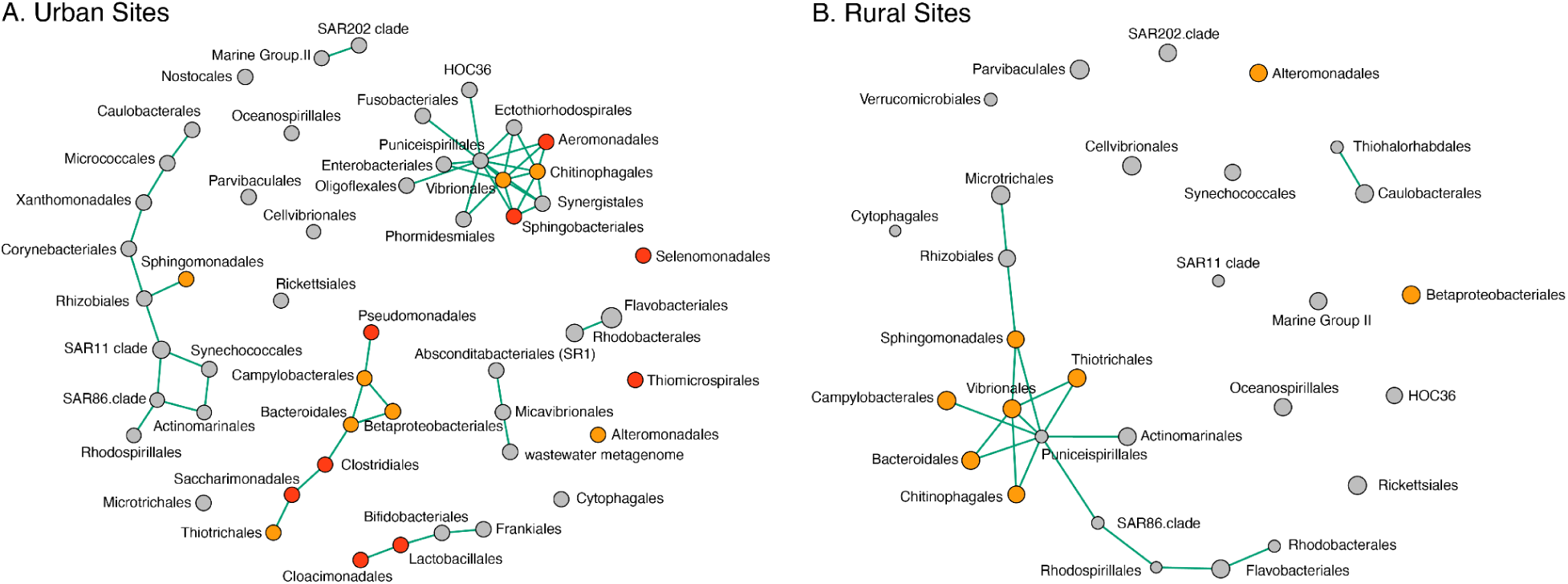
Covariance networks for bacterial orders detected at urban and rural sites. Orange nodes represent orders that are prevalent among samples from urban sites but were rarely observed in rural samples. Red nodes represent orders that are prevalent among or present in ≥ 3 samples from urban sites but not found in any rural samples. Green edges connecting nodes indicate significantly positive covariance between node orders (no negative relationships could be detected). There is a cluster composed solely of “red” (Pseudomonadales, Clostridiales, Saccharimonadales) and “orange” (Campylobacterales, Bacteroidales, Betaproteobacteriales, Thiotrichales) orders in the urban site network, indicating that these orders co-occur across samples collected from the urban sites

## Discussion

Coastlines around the world are rapidly being urbanized, with widespread consequences for adjacent marine ecosystems. Despite the apparent impacts of urbanization in many nearshore areas, it can be difficult to fully assess the effects due to shifting or absent baselines. Here, we specifically investigated the impacts of urbanization on the physicochemical conditions and microbial communities in nearshore marine ecosystems. Studying microbial responses to urbanization is a good first step in assessing the impacts of urbanization because changes in microbial communities can indicate disruption to the ecosystem at a larger scale (Nogales et al. 2011). Okinawa Island has undergone urbanization recently compared to many other coastal urban centers and the pattern of urbanization there allows for the resulting ecological effects to be investigated more directly. We detected an altered physicochemical setting in nearshore waters along urban coastlines compared to rural settings, with increased nutrient concentrations at urban sites (Fig. 2, 3). Microbial communities at urban sites were significantly different from communities at rural sites in all seasons (Fig. S4). The differences largely reflected increased microbial diversity at urban sites with the presence of bacterial taxa that were absent or rare at rural sites (Fig. 6, 7). Many of the additional taxa found at urban sites are associated with anthropogenic sources. Our results demonstrate that urbanization is changing the surrounding marine ecosystem and that mitigation approaches that include terrestrial management—such as restoring natural habitats along coastlines or improving wastewater treatment—may be more effective than approaches more focused on marine regulations.

### Urbanization alters the physicochemical characteristics of nearshore ecosystems

By monitoring physicochemical conditions at urban and rural sites along Okinawa Island, we documented a clear impact of urbanization on salinity and nutrient conditions in nearshore waters. Our results showed large variability in salinity at urban sites with surface water salinity dropping close to zero on several dates (Fig. 2). The salinity changes were most pronounced at the central and northern subsites, which were more likely to be directly impacted by the freshwater outlet of the adjacent watershed. The proximity to a freshwater outflow had a much less pronounced effect at the rural sites, despite the study watersheds all being of comparable size. The rural sites are semi-enclosed by intact coral reefs, which leads to longer residence times for inflowing freshwater and suspended sediments (Sakamaki et al. 2022). Longer residence times make it more likely for reduced salinity and increased nutrient-loading to be detected (Sakamaki et al. 2022). In contrast, the urban sites have undergone substantial land reclamation leading to the complete loss of the coral lagoon (Masucci and Reimer 2019), which should decrease the residence times for terrigenous inputs (Sakamaki et al. 2022). While the residence times for the specific locations included in this study are not yet determined, the projected variability in residence times based on site geomorphology do not match our observations. Thus, the difference in salinity between urban and rural sites is most likely due to the variation in impervious land cover (concretization) between sites. The urban watersheds in this study have very little natural or permeable ground cover remaining (Fig. 1), which increases flooding and run-off transported to coastal areas (Blum et al. 2020).

Wide variations in salinity—like those seen at the urban sites in this study—can have severe biological implications. Fluctuating salinity—especially reduced salinity from terrestrial run-off—reduces coral holobiont respiration and photosynthetic rates, heightens coral bleaching and mortality risk, disrupts symbiotic relationships between the coral, microbiome and zooxanthellae, and reduces coral larval and gamete survival (Röthig et al. 2023). As keystone organisms, coral mortality has downstream ecosystem and economic implications (Moberg and Folke 1999). Moreover, salinity is a major driver for bacterioplankton community composition (Lozupone and Knight 2007; Jurdzinski et al. 2023), and reduced salinity in nearshore regions can promote the growth of pathogenic bacteria (Bordalo et al. 2002; Randa et al. 2004; Burge et al. 2014) and increase the pathogenicity of some bacteria (Barca et al. 2023).

The major macronutrients—nitrogen and phosphorus—were enriched throughout the annual cycle at the urban sites compared to the rural sites in this study (Fig. 3). Both urban and rural land uses can contribute to nutrition loading. In urban areas, nitrogen and phosphorus pollution derive from sewage, fertilizers, detergents, and pet waste (Hobbie et al. 2017). In rural areas, nitrogen and phosphorus pollution derive mainly from agricultural fertilizers, manure application, and livestock waste (Del Rossi et al. 2023). However, nutrient export is positively correlated with impervious land cover, so while both urban and rural land uses produce excess nutrients, nutrients are more likely to reach coastal ecosystems in urban areas (Duan et al. 2012). While nitrogren and phosphorus were consistently elevated at urban sites compared to rural sites, they were highly variable (Fig. 3). Inorganic nutrients, such as nitrate + nitrite and phosphate measured in this study, are known to be temporally variable due to rapid uptake by microorganisms and short-term hydrodynamic processes (Viana and Bode 2013). It has, therefore, been recommended to measure dissolved organic carbon (DOM) or isotopic nitrogen signatures in macroalgae to reliably detect anthropogenic input to coastal ecosystems (Viana and Bode 2013; Sakamaki et al. 2022). However, by sampling inorganic nutrients over a time series in our study, we were able to detect increased nutrient loading at urban sites despite the variability. Moreover, our results largely align with those of Sakamaki et al. (2022), who chemically measured anthropogenic impact at a larger number of sites around Okinawa, but only at a single time point.

Nitrogen and phosphorus inputs to coastal waters are strongly connected to increased primary production (Howarth et al. 2021). Indeed, chlorophyll *a* fluorescence was significantly positively correlated with nitrogen and phosphorus concentrations at the urban sites we monitored (Fig. 4). Nutrient loading alongside stratification can also lead to hypoxia due to increased respiration and less mixing (Obenour et al. 2012), and dissolved oxygen was significantly negatively correlated with nitrate and nitrite at the urban sites in our study (Fig. 4). Nutrient loading is also detrimental to coral reefs. Increased nutrient availability fuels the growth of fleshy macroalgae that compete with coral for light and space and increases bioerosion by invertebrates (Silbiger et al. 2018). The negative effects of ocean acidification and rising temperatures on corals can also be amplified in high nutrient conditions (DeCarlo et al. 2015). Therefore, the altered near-shore nutrient conditions associated with urban land use on Okinawa are likely degrading the coral reef ecosystems that are important to the island for fisheries and tourism (Carlson 2015).

### Urbanization changes nearshore bacterioplankton community composition

We found significantly different bacterial communities in nearshore waters adjacent to urban areas compared to those adjacent to rural areas (Fig. 5, 6, 7). This difference in community composition is reflected in the elevated bacterial diversity found at urban sites (Fig. S3). In addition to a shared core bacterial community between the two site types made up of ubiquitous marine bacteria (Fig. 6), urban sites hosted additional taxa that were rare or absent at rural sites (Fig. 6, 7). Bacteria in the orders Cloacimonadales, Clostridiales, Lactobacillales, Pseudomonadales, Saccharimonadales, Selenomonadales, Sphingobacteriales, and Thiomicrospirales were present in at least three urban samples but absent in all rural samples (Fig. 6). Bacteria in the orders Aeromonadales, Alteromonadales, Bacteroidales, Betaproteobacteriales, Campylobacterales, Chitinophagales, Sphingomonadales, Thiotrichales, and Vibrionales were prevalent among urban samples but rare in rural samples (Fig. 6). Bacteria in many of these orders are known to be associated with anthropogenic sources, some are considered fecal indicator bacteria (FIB), and some are potentially pathogenic to humans and marine life (Verburg et al. 2021). In contrast, Verrucomicrobiales was the only order detected in multiple rural samples but absent in urban samples. Bacteria in the order Verrucomicrobiales are ubiquitous in soils (Bergmann et al. 2011), which may explain their absence in urbanized regions where there is little natural soil exposed (Fig. 1).

Among the bacterial groups that were present in urban regions but absent in rural areas (“red” orders, Fig. 6, 8), there are several of particular concern due to the inclusion of fecal indicator bacteria or known pathogens in the orders. The orders Clostridiales and Pseudomonadales were prevalent at both urban sites (Fig. 6, 7) and contain known human pathogens. When analyzed at the genus level, we found that most of the Clostridiales bacteria in our samples were classified as *Blautia spp.*, *Clostridium sensu stricto* 1, and several other genera in the Ruminococcaceae family. *Blautia spp.* and Ruminococcaceae bacteria are both common components of microbiomes in humans and other mammals, making up upwards of 50% of mammalian intestinal microbiome communities (Liu et al. 2021). *Clostridium sensu stricto* 1 includes the “true” *Clostridium* species, such as *Clostridium perfringens* (Yang et al. 2019). *C. perfringens* forms spores that are specific to sewage contamination and are stable in the environment. Thus, *C. perfringens* DNA detection reliably indicates human fecal contamination in environmental settings (Stelma 2018). The order Pseudomonadales includes several known pathogens in the genera *Pseudomonas* and *Acinetobacter*. However, Pseudomonadales ASVs in our dataset were classified as *Pseudomonas hussainii* and *Acinetobacter indicus,* which are not known human or animal pathogens. *P. hussainii* has been isolated from mangrove areas and, as a denitrifier, may be responding to increased nitrate concentrations at urban sites (Jaiswal et al. 2017). *A. indicus* was isolated from aquaculture waste for its chitinase activity and, in a reef ecosystem, may act as an invertebrate pathogen (Akram et al. 2022).

The orders Cloacimonadales, Lactobacillales, and Thiomocrospirales contain bacteria that are often associated with wastewater treatment plants, sewage sludge, and wastewater effluent (Meyer et al. 2016; Shakeri Yekta et al. 2019; Verburg et al. 2021). Bacteria in the order Cloacimonadales are members of sewage sludge bacterial communities and are associated with lipid and oil degradation (Shakeri Yekta et al. 2019). Bacteria in the order Lactobacillales are common components of human and animal microbiomes that, despite being non-spore-forming, often survive wastewater treatment and are released into surface water (Verburg et al. 2021). Sulfur-oxidizers, like those in the order Thiomicrospirales (Arora-Williams et al. 2022), are common in wastewater effluent (Meyer et al. 2016), can be indicative of low-oxygen conditions, and are capable of partial denitrification (Arora-Williams et al. 2022). High abundances of these microbes in urban ecosystems may be due to their presence in wastewater reaching the coastal ecosystem or they may be supported by nitrogen loading from wastewater or runoff.

Bacteria in orders that were prevalent in urban samples, but rare in rural samples (“orange” orders; Fig. 6, 8), also include groups of concern–such as Vibrionales, Bacteroidales, and Campylobacterales. The increased prevalence of Vibrionales bacteria at urban sites is alarming due to a recent surge in *Vibrio spp.* infections leading to human fatalities and limb amputations in developed coastal areas (Archer et al. 2023). Most severe vibriosis cases are caused by *V. vulnificans*, but multiple *Vibrio* species can cause serious illness in humans (Baker-Austin et al. 2018) and marine organisms (Grimes 2020). Here, we found that *Vibrio fortis* and *Vibrio chagasii* were abundant in samples from U1 - Ginowan, whereas *Photobacterium leiognathi* subsp. *mandapamensis* was the major Vibrionales species found at U2 - Nago, and *Photobacterium damselae* subsp. *damselae* was detected at both sites. *V. fortis* and *V. chagasii* are both marine pathogens—*V. fortis* causes coral bleaching (Sun et al. 2023) and enteritis in fishes (Wang et al. 2016) while *V. chagasii* infects molluscs (Urtubia et al. 2023). *P. mandapamensis* is a bioluminescent symbiont of several reef fishes common in coastal water around Okinawa and can also be found free-living, especially where host fish are abundant (Urbanczyk et al. 2011). *P. damselae* is a pathogen of marine animals–including vertebrates and invertebrates–as well as in humans, where it causes necrotizing fasciitis (Rivas et al. 2013). Given that the Vibrionales bacteria detected here are marine, their increased abundance at urban sites could be due to changing conditions in the nearshore ecosystem caused by urbanization rather than direct anthropogenic sources. For instance, several marine *Vibrio* species have previously been shown to become more prevalent and infectious when salinity is reduced to 5–25 PSU (Sullivan and Neigel 2018) and we frequently recorded values within this range at urban sites in our study (Fig. 2).

Bacteria in the orders Campylobacterales, Betaproteobacteriales, and Bacteroidales were the most abundant groups that were prevalent among urban samples but not rural samples (“orange” orders; Fig. 6, 7). Campylobacterales bacteria are clinical pathogens (Man 2011), but can also be environmental bacteria (Fera et al. 2004), and pertinent to our study, are abundant in urban road-run-off samples (Liguori et al. 2021). The majority of Campylobacterales ASVs in our study were classified as *Arcobacter spp*. Of the five species that comprise the genus *Arcobacter*, three are associated with human disease and are found in human diarrhea and sewage samples (Fera et al. 2004). However, *Arcobacter spp*. are also found in a variety of environmental settings—including rivers, salt marshes, and nearshore marine waters—indicating that water may be a major transmission route for *Arcobacter* infection (Fera et al. 2004). Betaproteobacteriales is a large and diverse taxonomic group, but many members of this group are enriched in samples from wastewater treatment plants (Zhang et al. 2017). Betaproteobacteriales ASVs in our samples belonged to 102 separate genera, but C39 (family: Rhodocyclaceae) was by far the most abundant, and is associated with pollution and wastewater input in coastal ecosystems (Nascimento et al. 2018; Kopprio et al. 2020). The Bacteroidales order includes bacteria that make up the majority of mammalian gut microbiomes (Magne et al. 2020), are members of sewage sludge communities (Yekta et al. 2019), and have been identified in road run-off (Liguori et al. 2021). In our study, the majority of Bacteroidales ASVs abundant at urban sites belonged to the genera *Bacteroides* and *Prevotella*, which are major mammalian gut bacteria and likely originate from sewage (Wu et al. 2011).

Overall, our results showcase a clear anthropogenic impact on microbial communities in nearshore environments adjacent to heavily urbanized watersheds in Okinawa. The urban ecosystems were consistently altered rather than exhibiting episodic or seasonal disturbances. Previous work focusing on the Tancha (R1) sampling site showed that extreme run-off events associated with typhoons caused short-lived perturbation of nearshore microbial communities, with the bacterial community composition returning to baseline only 48–72 hours after storms passed (Ares et al. 2020). The quick return to pre-typhoon conditions highlights the quick-flushing of Okinawa’s coral lagoons, as well as the overall resiliency of nearshore ecosystems in more rural regions of Okinawa’s coast. In contrast, the consistently altered state observed at urban sites may indicate that these ecosystems are so altered that they have reached a tipping point and experienced a regime change. This regime change may interact with other components of the system (e.g., corals, algae, and other invertebrates) and create feedback loops that further degrade the ecosystem and its functioning (Qin et al. 2020; Becker et al. 2023). Alternatively, the consistently altered microbial communities observed at urban sites may reflect sustained, ongoing disturbances that would likely also have downstream ecological consequences (Ruprecht et al. 2021).

### Conclusions & Future Directions

Biweekly observations made in nearshore waters along a large urban-to-rural gradient revealed the profound influence of urbanization on salinity, macronutrient concentrations, and microbial communities in a recently urbanized subtropical island system. Expanding this work in the future to include metatranscriptomics would allow for better characterization of the active metabolisms of microbes in urban and rural areas and could shed light on microbial nutrient uptake, as well as pathogenicity. Nonetheless, the observed changes in physicochemical parameters and microbial communities have likely contributed to degrading nearby coral reefs (Hasegawa 2011; Heery et al. 2018), as coral disease, death, and loss of diversity have been attributed to reduced salinity, nutrient loading, and introduced pathogenic bacteria in other systems (Carlson et al. 2019). Coral reefs around Okinawa and other subtropical and tropical islands provide myriad ecosystem services, including supporting tourism and fisheries as well as protecting islands from waves and erosion, but the ability of reefs to perform these ecosystem services is declining (Eddy et al. 2021). To protect these key cultural, economic, and ecological resources, land use and coastal development need to proceed mindfully and a “ridge-to-reef” management mindset should be adopted. Ridge-to-reef management emphasizes the land-sea connection and advocates for restoring natural habitats—such as forests, wetlands, seagrass meadows, and mangroves—or practicing more sustainable agriculture on coastlines (Carlson et al. 2019). Islands, such as Okinawa, have limited geographic area available, and as a result, development often abuts the coast. Adding back natural buffers between urban development (or intensive agriculture) and the coast can reduce freshwater input, absorb nutrients, catch sediment, and filter out pollutants (Carlson et al. 2019). Moreover, more comprehensive sewage treatment and moving outfalls further offshore can significantly reduce nutrient and pathogen loading on reefs (Bahr et al. 2015). Such mitigation efforts may become more critical as cyclonic tropical storms increase in both frequency and intensity due to global climate change (Li and Chakraborty 2020). Tropical cyclones deliver large amounts of precipitation to Okinawa that trigger extreme runoff events (Ares et al. 2020). Rural sites in Okinawa have proven to be resilient to extreme storms (Ares et al. 2020), but more study is needed to understand how such events may differentially impact the more disturbed urban ecosystems identified in this study.

## Acknowledgments

We acknowledge Okikanka for collecting samples, and Ayse Oshima and Pradeep Palanichamy for assisting in filtering water samples and extracting DNA. The Okinawa Institute of Science and Technology (OIST) Sequencing Center (SQC) performed sequencing library preparation and sequencing. Research was funded by the OIST Marine Biophysics Unit and a Japan Society for the Promotion for Science (JSPS) Kakenhi award to AA (award 20K19986). MMB was supported by a Simons Foundation Postdoctoral Fellowship in Marine Microbiology (award 874439).

## Author Contributions

AA conceived of the project. AA, SM, and MMB planned research. AA and YY performed research. AA, MMB, and KD analyzed data. MMB wrote the manuscript. All authors edited drafts and approved the final version of the manuscript.

## Data availability statement

All raw sequencing reads generated for this project are publicly available from November 23rd of 2023 with accession number PRJNA1044524. Intermediate data files (including CTD and nutrient measurements) and the code necessary to replicate analyses are available in a GitHub repository (https://github.com/maggimars/UrbanOki) and as an interactive HTML document (https://maggimars.github.io/UrbanOki/Amplicons.html).

