## Supplementary Material for "Urbanization of a subtropical island (Okinawa, Japan) alters physicochemical characteristics and disrupts microbial community dynamics in nearshore ecosystems"

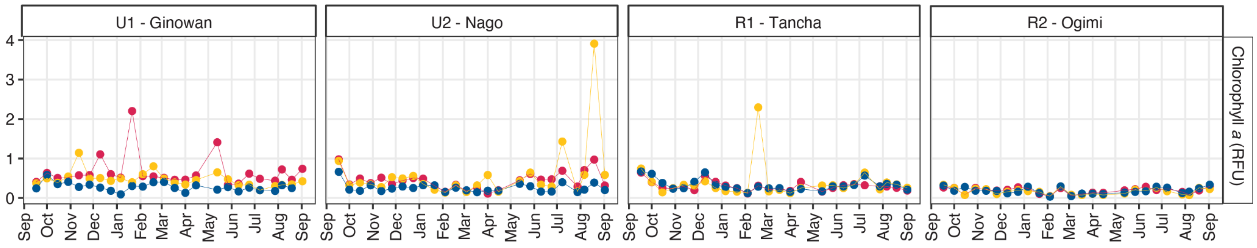


**Fig. S1** Time series of Chlorophyll *a* fluorescence measured at nearshore sites in urban and rural areas along the west coast of Okinawa Island, Japan. Chlorophyll *a* fluorescence was measured at each subsite (south, central, and north) within each sampling site (U1, U2, R1, and R2) using a RINKO CTD probe. Plots are faceted by sampling site. Point and line color represents subsites—blue, south (S); red, central (C); yellow, north (N)—where water samples and measurements were taken. Chl *a* was elevated at the urban sites compared to rural sites, especially at the central (red) and north (yellow) subsites

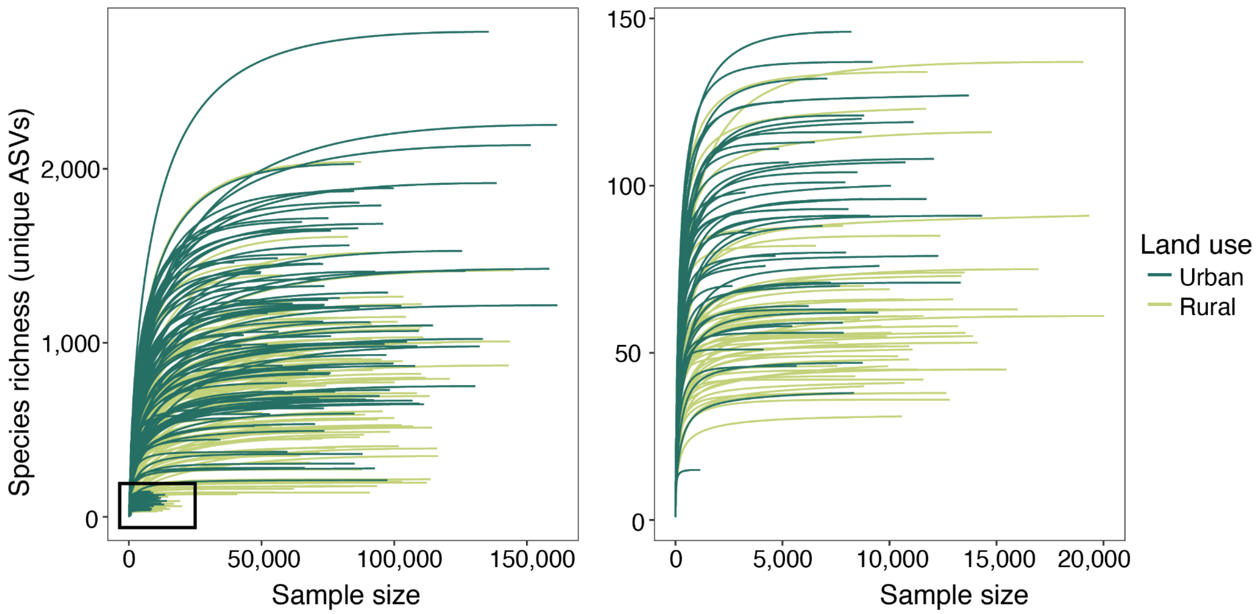


**Fig. S2** Rarefaction Curves for all samples from urban and rural sites across the time series. Rarefaction was performed with the phyloseq R package and results were plotted with the `ggrare()` function. Line color represents the sampling area type (Urban: dark green, Rural: light green). The plot on the right is a zoomed-in view of the left plot area highlighted by the black box. Samples with low observed species richness saturate before reaching the maximum sequence sample size. However, the sample with the lowest species richness was removed from the dataset before further analysis due to the very low number of total sequences

A.

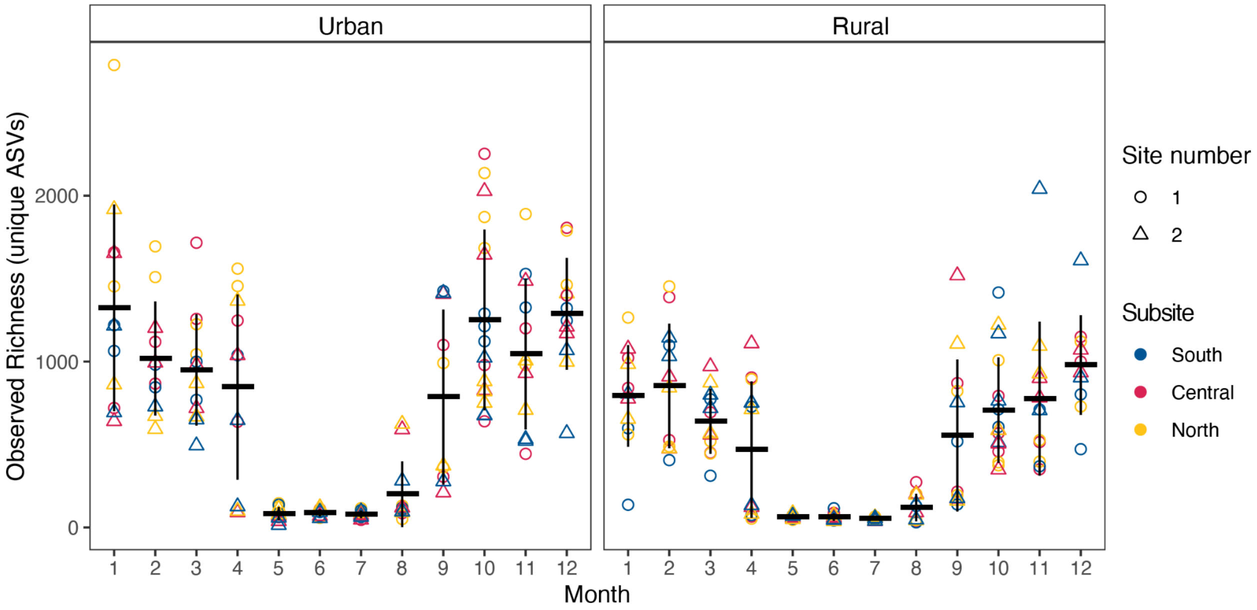


B.

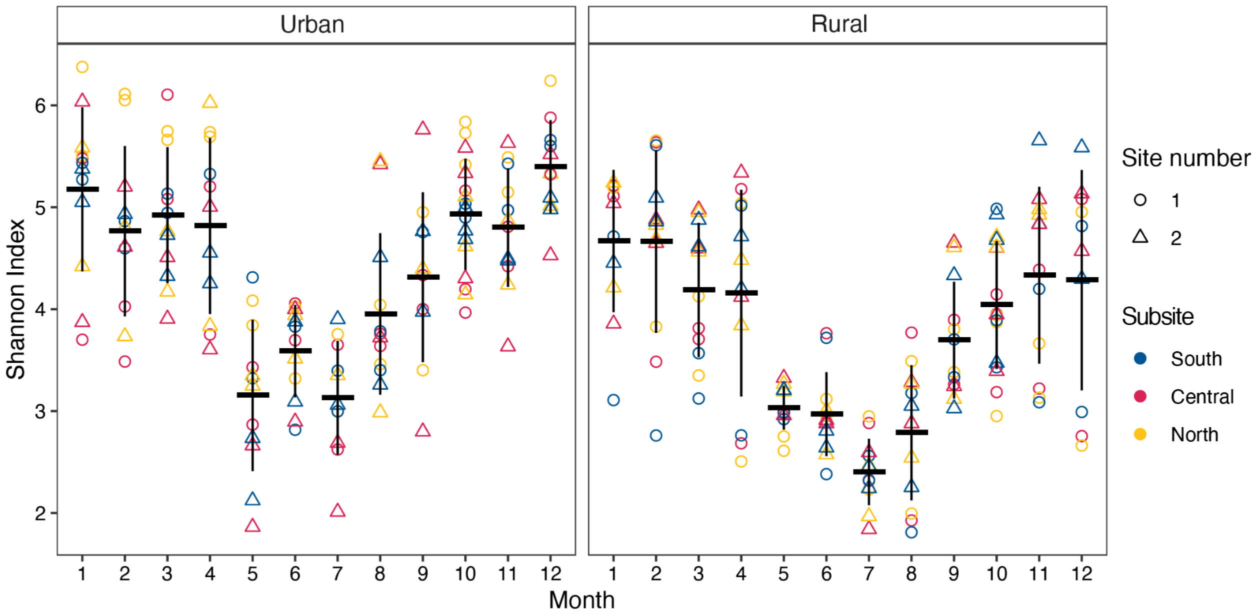


**Fig. S3** Alpha diversity indices for nearshore bacterial communities evaluated at urban and rural sites along Okinawa Island. (A) Observed richness (number of unique ASVs) in each sample represents total diversity, whereas (B) the Shannon Index is a metric that combines total diversity with the evenness of ASV abundance in each sample. Alpha diversity estimates (richness and Shannon indices) were determined with the 'breakaway' R package. Urban sites had elevated alpha diversity (richness and Shannon indices) throughout the seasonal cycle. Richness and Shannon indices were lower during the summer months than the rest of the year at all sites

| **Month** | **Richness (unique ASVs)** | **Shannon Index** |
|---|---|---|
| January | * (*p*=0.027) | * (*p*=0.037) |
| February | *n.s.* (*p*=0.298) | *n.s.* (*p*=1.0) |
| March | * (*p*=0.021) | * (*p*=0.024) |
| April | *n.s.* (*p*=0.079) | *n.s.* (*p*=0.190) |
| May | *n.s.* (*p*=0.078) | *n.s.* (*p*=0.478) |
| June | * (*p*=0.008) | * (*p*=0.004) |
| July | * (*p*=0.004) | * (*p*=0.001) |
| August | *n.s.* (*p*=0.355) | * (*p*=0.001) |
| September | *n.s.* (*p*=0.169) | * (*p*=0.036) |
| October | * (*p*=0.001) | * (*p*=0.000) |
| November | *n.s.* (*p*=0.078) | *n.s.* (*p*=0.242) |
| December | * (*p*=0.02) | * (*p*=0.003) |

**Table S1.** Statistical significance of differences in alpha diversity indices based on sampling area type (i.e., urban or rural). Pairwise Wilcox tests were performed for each month of the sampling year and results were considered statistically significant if the *p*-value was less than or equal to 0.05. Statistically significant results are indicated with an asterisk ('*') and nonsignificant results are indicated '*n.s.*'.The difference in observed richness between urban and rural sites was statistically significant for 6 out of 12 months and the difference in the Shannon index was statistically significant for 8 out of 12 months

A.

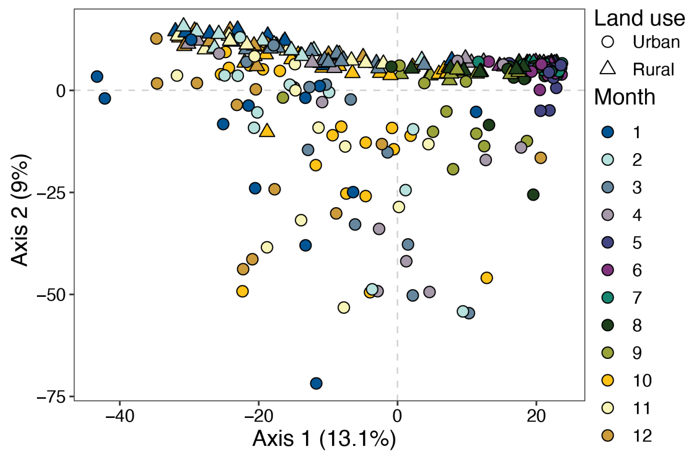


B.

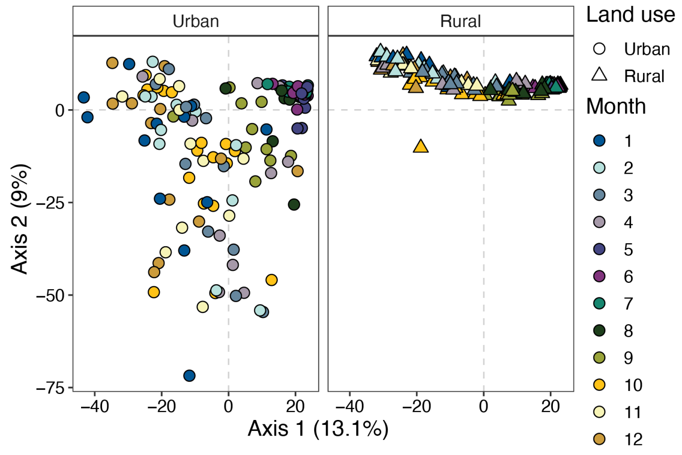


**Fig. S4**. Principal coordinate analysis (PCoA) plots of Aitchison distances between bacterial communities through time at two near-shore sites adjacent to rural areas and two near-shore sites adjacent to urban areas. (A) is a PCoA ordination plot of all samples and (B) is the same ordination but urban and rural samples are plotted on separate axes to better show the separation between sample types. Shape indicates sampling sites, with circles representing urban sites and triangles representing rural sites. The point color indicates the month of the year, with blues representing winter months, purples representing spring months, greens representing summer, and yellows representing autumn. Samples from spring and early summer cluster together (top right quadrant) regardless of site type. Urban sites show much more spread overall and urban samples from the rest of the year do not cluster with rural samples. A permutational multivariate analysis of variance (PERMANOVA) with 999 permutations indicated that bacterial communities were significantly different based on sample site land use (urban or rural; $R^2$=0.03, $F$=8.54, $p$=0.0011)

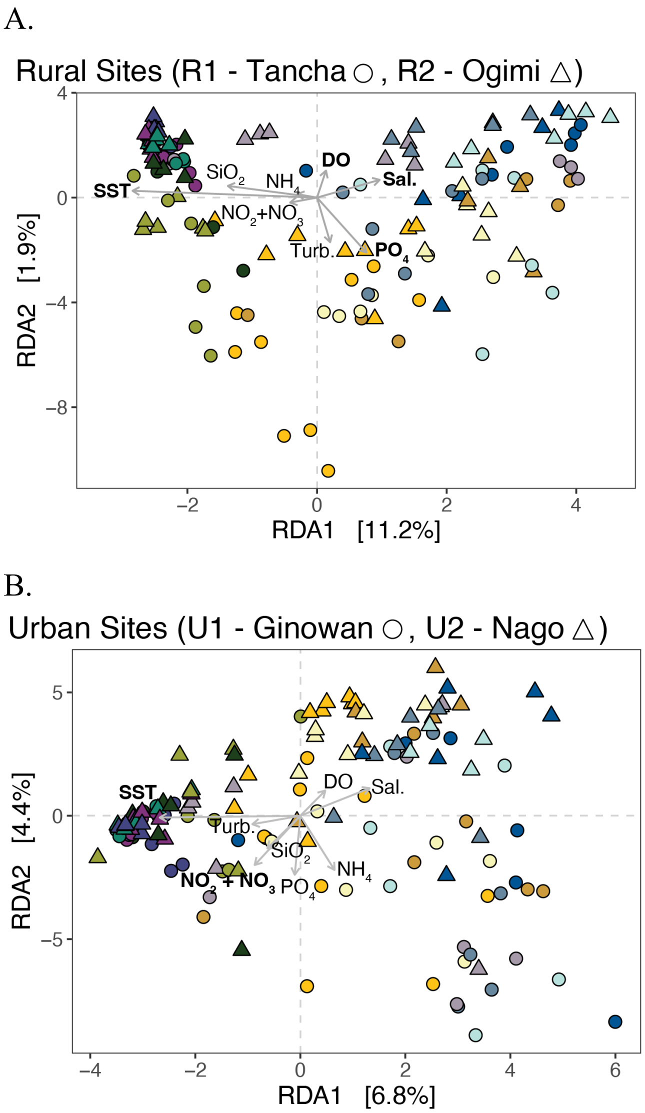


**Fig. S5.** Redundancy analysis (RDA) based on Aitchison distances between bacterial community samples and z-scored environmental variables for rural sites (A) and urban sites (B). Redundancy analyses were performed with the ‘vegan’ R package. Analyses of variance (ANOVA) performed on RDA results indicated that the RDA models were statistically significant for both rural (999 permutations, $F$=4.04, $p$=0.001) and urban sites (999 permutations, $F$=2.28, $p$=0.004). ANOVAs by term were subsequently run to determine which variables were significantly contributing the the RDA results (Table S2). Variables with a $p$-value $\leq$ 0.05 were considered statistically significant and highlighted on the plot by bolding the variable name. In the RDA for rural samples (A), SST, DO, Salinity, and $PO_4$ were statistically significant. In the RDA for urban samples (B), SST and $NO_2$+$NO_3$ were statistically significant. A color-blind accessible version of this plot is available at https://maggimars.github.io/UrbanOki/Amplicons.html#with_colorblind_colors59

A. Rural

| **Variable** | ***F* (Term RDA)** | ***p* (Term RDA)** | ***$R^2$* (Variance partition)** |
|---|---|---|---|
| $NO_2$+$NO_3$ | 0.64 | 0.571 | |
| $NH_4$ | 0.17 | 0.959 | |
| $PO_4$ | 6.9 | 0.001 * | 0.038 |
| $SiO_2$ | 2.2 | 0.088 | |
| SST | 4.2 | 0.008 * | 0.028 |
| Salinity | 2.8 | 0.04 * | 0.013 |
| Turbidity | 2.0 | 0.116 | |
| DO | 13.4 | 0.001 * | 0.080 |

B. Urban

| **Variable** | ***F* (Term RDA)** | ***p* (Term RDA)** | ***$R^2$* (Variance partition)** |
|---|---|---|---|
| $NO_2$+$NO_3$ | 5.53 | 0.002 * | 0.035 |
| $NH_4$ | 0.77 | 0.455 | |
| $PO_4$ | 1.48 | 0.156 | |
| $SiO_2$ | 1.06 | 0.356 | |
| SST | 3.76 | 0.004 * | 0.028 |
| Salinity | 1.85 | 0.095 | |
| Turbidity | 2.29 | 0.061 | |
| DO | 1.49 | 0.162 | |

**Table S2.** Analysis of Variance (ANOVA) by term results for redundancy analysis (RDA) models for rural (A) and urban (B) sites based on Aitchison distances for microbial communities and z-scored environmental variables. When term results were statistically significant ($p \leq 0.05$, marked by an asterisk ('*')), variance partitioning was performed for that term and the results are included in the table

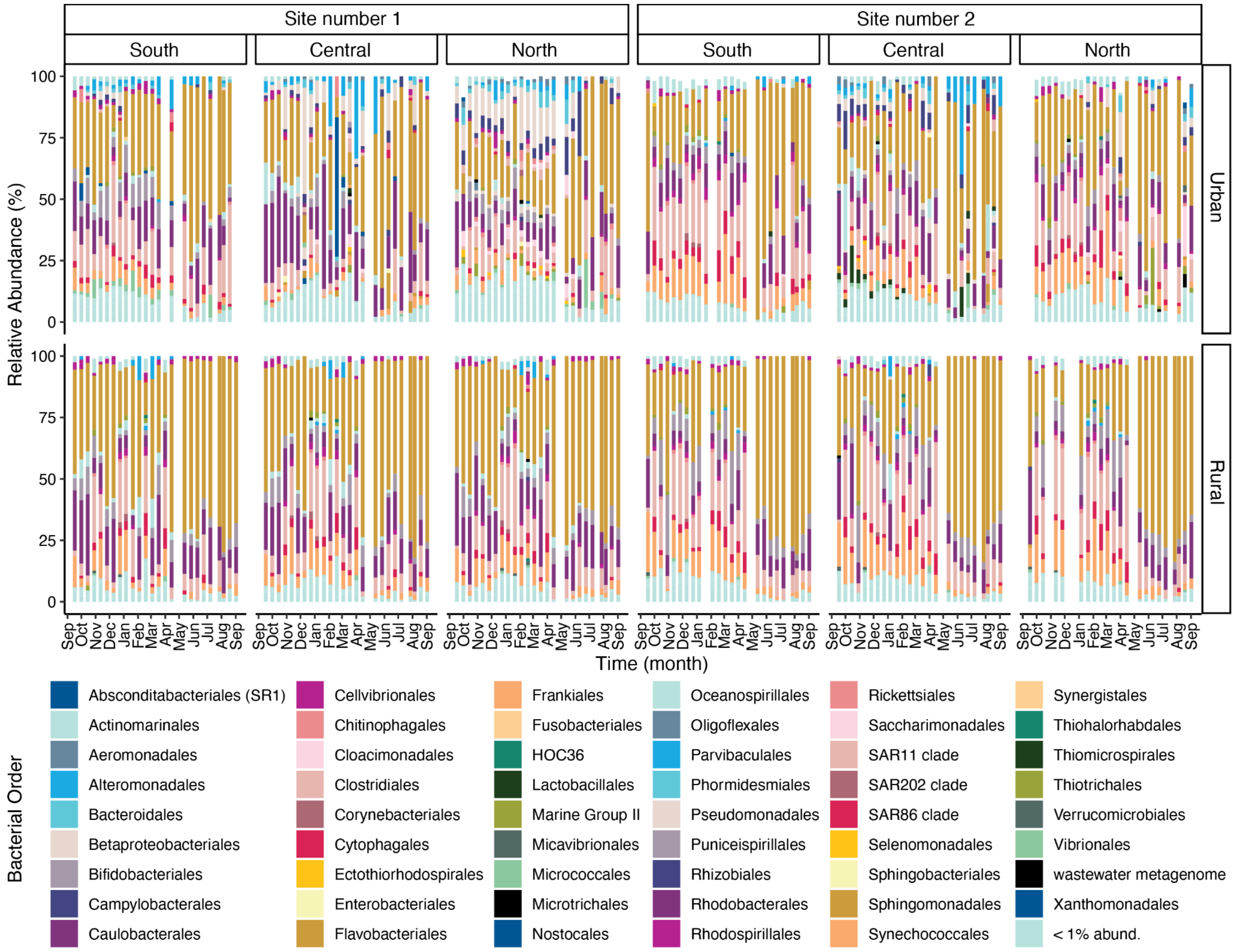


**Fig. S6** Relative abundance of bacterial orders across a time series collected at urban and rural sites along the west coast of Okinawa Island. The plot is faceted by sampling area, with urban and rural sites separated vertically and samples grouped by site number and subsite (S; southern subsite, C; central subsite, and N; northern subsite) horizontally. The stacked bar chart fill colors indicate bacterial order at each sampling date (twice a month). The order Flavobacteriales is most abundant across the time series at all sites and subsites. Synechococus, SAR11, and Rhodobacterales are also abundant across all samples. Finer scale variability is better visualized in Figures 6 and 7 in the main text
